# Experimental study of social learning in three wild shell-dwelling Tanganyikan cichlids that vary in sociality

**DOI:** 10.1101/2025.03.06.641821

**Authors:** Maëlan Tomasek, Valérie Dufour, Alex Jordan

**Author notes:** These authors contributed equally (Valérie Dufour, Alex Jordan). **Corresponding author: Maëlan Tomasek**, Social and Cognitive Ethology Team, UMR 6024, LAPSCO, BAT 40, Campus CNRS, 23 rue de Loess, 67037 Strasbourg, France.

## Abstract

The social intelligence hypothesis posits that animals living in more complex social groups display better cognitive performances. However, this hypothesis has mainly been investigated in primates and studies using similar paradigms across different species are scarce. We tested three species of wild Lamprologine shell-dwelling cichlids from Lake Tanganyika (*Neolamprologus multifasciatus*, *Lamprologus ocellatus*, and *Lamprologus ornatipinnis*) that vary in their levels of sociality, in identical colour associative learning and social learning tasks. We found species differences in engagement in the training for these tasks and only 5 out of 24 individuals learnt to feed from a neutral object (a shell). One *L. ocellatus* and one *L. ornatipinnis* learnt to choose the correct colour in the associative learning task and were further used as demonstrators in the social learning task. In this task, no species showed signs of colour stimulus enhancement as the observers’ choices of colours were not influenced by the demonstrators. However, we found evidence for stimulus enhancement at a larger scale as naïve observers approached the novel experimental apparatus faster than the demonstrators when first exposed to this apparatus. Our results encourage further research of social learning processes in individual fishes and highlights the difficulties in using identical paradigms in comparative psychology, even across very close species.

## 1. Introduction

Many hypotheses have been put forward to explain the evolution of cognitive abilities in animals (Henke-von der Malsburg et al., 2020). Coping with the challenges of living in a social group is one of them, known as the social intelligence hypothesis (Dunbar, 1998). According to this hypothesis, animals living in social groups should evolve bigger and more complex brains as well as greater cognitive abilities (Dunbar, 2009), especially in the context of social cognition. Indeed, species living in social groups should have access to more information from their conspecifics than non-grouping species and should therefore have greater opportunity for social learning (Danchin et al., 2004), defined as “any learning process resulting from the behaviour of other animals” (Shettleworth, 2010). Indeed, beyond the information acquired via personal experience (private information), animals can access diverse information like predator location, new foraging areas, or handling techniques by watching con- or heterospecifics (public information) (Shettleworth, 2010;Laland, 2004; Avarguès-Weber et al., 2013). Several mechanisms of social learning exist, from simple stimulus enhancement (increased likelihood of contacting a type of stimulus by observing others doing it) or local enhancement (increased likelihood of visiting a place by observing others going there), to more complex forms of emulation or imitation (Shettleworth, 2010). To assess social learning in animals, we can thus test whether the observation of a demonstrator performing a task influences the later performance of the observer in the same task (*e.g.,* Dindo 2010, 2011).

The social intelligence hypothesis has been developed and mainly tested in primates but requires testing in a broader range of taxa to gain a more thorough evolutionary and ecological basis. To properly test the hypothesis, species studied should be phylogenetically close as differences in cognitive abilities could be attributed to phylogenetic signal or differences in perceptual, physiological, or morphological traits (Bräuer et al., 2020; MacLean et al., 2012). In this respect, cichlid fishes represent an ideal system. Cichlids are a model of adaptive radiation, especially in the African Great Lakes, and their phylogeny is thus well studied and established (Matschiner et al., 2020; Ronco et al., 2021; Salzburger, 2018). Moreover, while phylogenetically close, they present a wide range of socio-ecological attributes which makes them ideal candidates to investigate hypotheses about the evolution of cognition (Lein & Jordan 2020). While it is known that fishes broadly possess the capacity for social learning (Brown & Laland, 2003), it is not yet clear whether more social species are better at it than others. For instance, non-grouping teleost species watching heterospecifics have been shown to have the same local enhancement aptitudes than grouping species (Webster & Laland, 2017). In cichlids, some studies have compared the cognitive abilities of species differing in their social organisation in the Lamprologini tribe of Tanganyikan cichlids. In this tribe, the more social *Neolamprologus pulcher* performed better at an operant conditioning task than less social *Neolamprologus caudopunctatus* (Fischer et al., 2021). However, in other learning tasks, non-social species have performed better than social ones (Stanbrook et al., 2020b). Most social learning studies in cichlids have shown social facilitation mechanisms (increased likelihood to perform a behaviour when in the company of others performing it, Shettleworth, 2010) in operant conditioning tasks with groups of two or more individuals (Karplus et al., 2007 in *Oreochromis niloticus*, Zion et al., 2011 in *Sarotherodon galilaeus*, Rodriguez-Santiano et al, 2020, 2022 in *Astatotilapia burtoni*, Long & Fu, 2022 in *Chindongo demasoni*, and Stanbrook et al., 2020 in Lamprologine cichlids). However, to the best of our knowledge, only one study has investigated social learning mechanisms at the individual scale and found no evidence for observational conditioning (associating a cue or object with an affective state or behaviour by watching demonstrators respond to it, Shettleworth, 2010). In the task, South American convict cichlids *Amatitlania siquia* could observe a demonstrator showing an avoidance response to a novel object. However, when presented with the same object, the observers did not show the same avoidance response (Barks & Godin, 2013). It is therefore not yet clear whether the social intelligence hypothesis is supported in cichlids. Moreover, to properly test this hypothesis and yield comparable results, it would be necessary to develop a standardized paradigm that could be used reliably in different species.

Here we studied three species of wild Lamprologine cichlids in an associative learning and a social learning task: *Neolamprologus multifasciatus*, *Lamprologus ocellatus* and *Lamprologus ornatipinnis*. All three species lineages are phylogenetically close as they diverged approximately 4 million years ago (Ronco et al., 2021). Moreover, as many other shell-dwelling species of this tribe, they present different levels of sociality while living in the same environment (Lein & Jordan, 2020). *N. multifasciatus* is a social species living in groups of more than ten individuals in a small territory (Bose et al 2022). *L. ocellatus* has a more intermediate level of sociality and lives mostly solitarily or in pairs or harems of several individuals. *L. ornatipinnis* is a solitary species living in a wide territory. While one previous study has trained captive-bred *N. multifasciatus* to push a lid to uncover a hole containing a food reward (Stanbrook et al., 2020a), ours is the first study to use *L. ocellatus* or *L. ornatipinnis* in associative and social learning contexts, and the first study of its kind to use wild subjects in any of these species. We aim to develop a new paradigm to test associative learning and social learning abilities in these species and explore potential interspecific differences. According to the social intelligence hypothesis, we predict that *N. multifasciatus*, the more social species, should display better cognitive abilities than the others and should therefore perform better at the tasks, especially in the social learning task. The performance of *L. ocellatus* should be poorer than *N. multifasciatus*, but better than *L. ornatipinnis,* which has the least complex social organisations.

## 2. Material and methods

### Subjects and housing

The fishes used in the experiments were collected in the wild in Zambian parts of Lake Tanganyika from April 14^th^ to 16^th^, 2022 (in front of Chikonde village (8°71’49.2”S, 31°12’53.1”E) for *N. multifasciatus* and *L. ornatipinnis* and in Isanga bay (8°64’99.3”S, 31°19’10.7E) for *L. ocellatus*). Eight individuals of each species were taken with the shell they lived in and were brought to outdoor ponds filled with lake water. All fishes were adult males, though sex determination could not be entirely assured without dissection, which was not carried out. An entry clinical exam by a trained veterinarian (MT) revealed no important health anomalies in our fishes apart from occasional scraping behaviour indicating mild external parasitosis in some individuals.

The fishes were housed in outdoors 95×80×50cm (∼320L) concrete tanks layered with sand taken from the lake. They were kept from April 14^th^ to May 14^th^, 2022, for the whole duration of the experiments. Individuals were grouped by pairs, and each pair was given a home tank. Pairs were composed by selecting two size-matched fishes. Their home shells were placed in each tank at distances adapted to each species (farther apart for *L. ocellatus* and *L. ornatipinnis* to avoid any aggression but closer for *N. multifasciatus* to maintain social interactions). A 50% water change was conducted weekly with fresh water pumped from the lake, and fishes were housed at a density of 1 fish per 160L water. In addition to the food provided during the experiments, the animals could feed *ad libitum* from the small invertebrates contained in the lake water, which is their natural diet (Lein & Jordan, 2021). At the end of the experiments, fish were returned to their collection sites in the lake.

Several food rewards were tested to find the most suitable and attractive one for wild fishes. The only food rewards for which they appeared to be motivated was live *Artemia* brine shrimp that were hatched at the onshore laboratory. Although ensuring motivation of the fishes, these small size and live shrimp constituted a reward which was not granular, contrary to standard food rewards used with captive-bred fishes. Instead, the shrimp diffused rapidly in the water and the apparatuses needed to be designed in order to contain them and avoid this diffusion. Fishes did not show any interest in more standard food rewards.

### Training

The training took place from April 19^th^ to May 1^st^, 2022. Individuals were trained to feed from a 3D-printed white replica *Neothauma* shell in which brine shrimp were injected via opaque tubing. The training was divided in three stages: stage 1 consisted in learning to eat brine shrimp from a tube connected to a syringe, stage 2 consisted in learning to eat the shrimp coming from that tube in an open white 3D-printed semi-shell, and stage 3 consisted in learning to go eat from the complete shell (**Figure 1A**). An individual was deemed ready to pass to the next training stage once it did not show any signs of avoidance toward the apparatus but readily approached it and fed. We conducted two training sessions per day, in the morning and in the evening, as the fishes were found to be less active during midday. Individuals were not separated during the training and sometimes both individuals could come eat to the shell together. We subsequently separated the individuals that had succeeded in stage 3 of training into “demonstrators” and “observers” in the next experiments.

**Figure 1:**
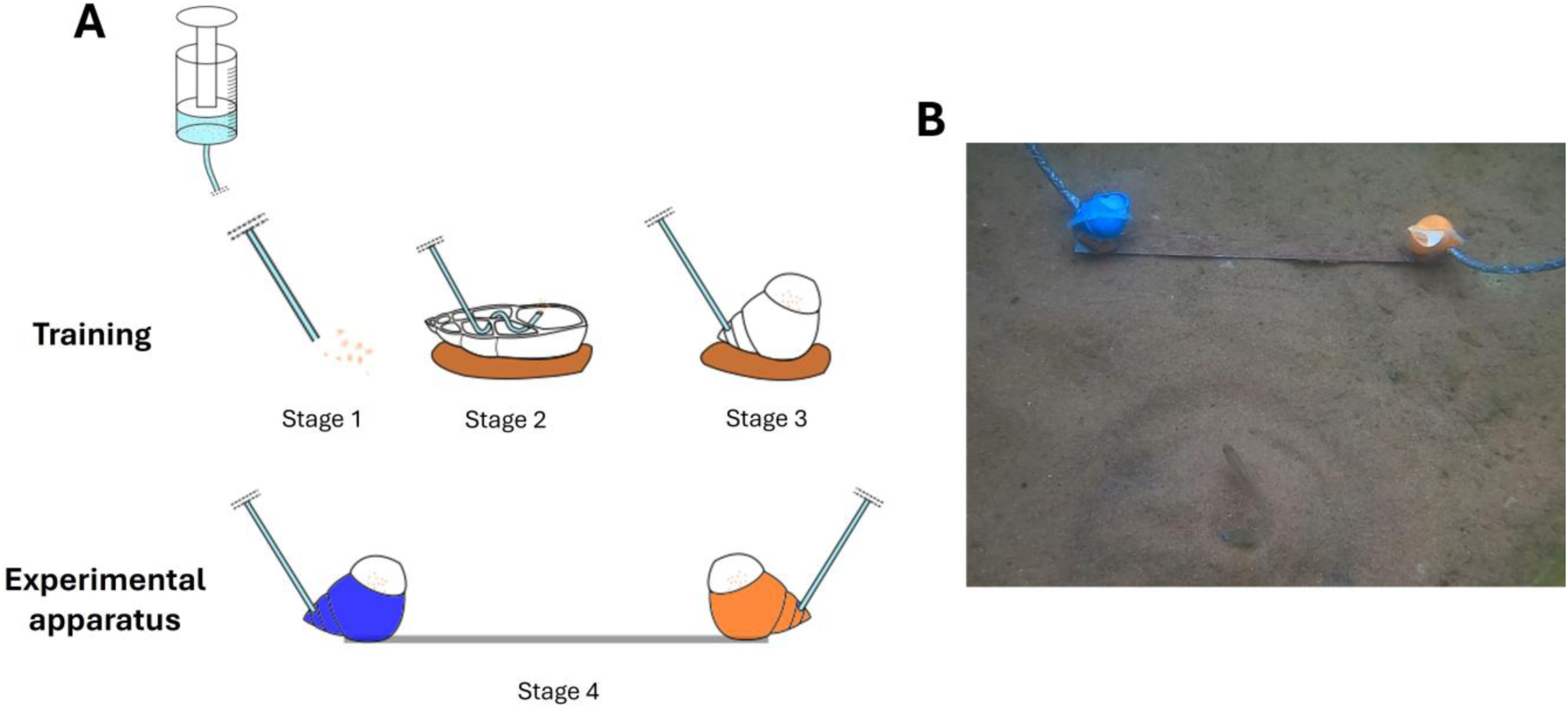
Learning process of the associative learning paradigm. **A.** The different apparatus for each learning process stage, from training to associative learning experiments. *Artemia* brine shrimp were injected with a syringe in the tubes. **B.** Image of a *L. ocellatus* (Mac) at his home shell prior to engagement with the experimental apparatus.

### Testing associative learning for demonstrators

This experiment was conducted after the training stages from May 2^nd^ to May 6^th^, 2022. In this task, demonstrators had to learn to feed in either an orange or a blue shell when given the choice between both colours (stage 4 apparatus, **Figure 1**). For each trial, the experimenter waited for the focal fish to be located at its home shell before presenting the apparatus, to ensure that both coloured shells were at equal distance from the starting point of the fish. When the individual was presented with the apparatus, it could choose to go to the orange or the blue shell. Only one colour was rewarded (assigned randomly across individuals). If the fish went to its correct shell (defined as the fish’s head passing the threshold of one of the shells), brine shrimp were injected in the shell and the fish could feed for several seconds. In the first six sessions, if the fish went to the incorrect shell, the shell was simply unrewarded, and the fish could still go to the correct shell and get rewarded. In the subsequent sessions, if the fish went to the incorrect shell, the shell was unrewarded, the apparatus was removed, and the trial was ended. If during a trial, a fish did not show any interest in the apparatus for at least 5 minutes, the trial was ended (“no choice”) and a new trial began after a short pause of a few seconds.

We conducted ten sessions of the associative learning task. In each session, the fish was presented between five to eight times with the apparatus depending on its motivation to investigate the shells. Indeed, it could occur that brine shrimp escaped the apparatus after several trials. The fish would then just eat them from the water column and would not pay attention to the apparatus anymore. The location of the rewarded shell (to the right or the left of the home shell) was randomized throughout trials to promote colour and not location learning. The rewarded shell was at the same location a maximum of two times before switching. During all sessions, the future “observer” fish which was housed in the same tank and was placed under an opaque container to avoid any visual contact with the task and the tested fish.

Cichlids are tetrachromatic and are therefore supposed to distinguish orange and blue (Sabbah et al., 2010). No specific visual study on our species of interest has ever been conducted but some other Lamprologine fishes were found to be able to perceive those colours (Egger et al., 2011).

### Testing social learning in naïve observers

This experiment was conducted from May 6^th^ to 9^th^, 2022. In this task, naïve observers, which had never been exposed to the two-coloured shell apparatus, could first observe a demonstrator going to feed in one specific colour of this apparatus. The observers were then tested with the same apparatus to see whether their choices were influenced by this observation. First, for three sessions, the observer fish was placed under a transparent container and could observe the demonstrator feeding from the coloured shell apparatus. The demonstrator only fed in one of the coloured shells. As before, if the demonstrator approached the wrong colour, the apparatus was removed without feeding. After each observation session, the observer could then feed in the neutral white stage 3 shell as usual. Second, for the next four sessions, the observer could first observe the demonstrator feeding from a specific colour five trials in a row and only then was presented with the two-coloured shell apparatus between five to seven times depending on their motivation. Both shells were rewarded for the observer to avoid private learning experience and to nurture their interest in the task. When an observer chose a shell (whatever its colour), it could take five mouthfuls before the apparatus was removed to ensure equal potential experience for both colours. The location of the shells was also randomized for the observer as described earlier. If fishes show social learning, more particularly stimulus enhancement towards colour, then the observer should go feed more frequently in the coloured shell that had been demonstrated.

### Testing social learning in experienced observers

This experiment took place from May 11^th^ to 14^th^, 2022. In this task, experienced observers (*i.e.*, former demonstrators), which had previously learnt to go to a specific colour, could first observe a demonstrator going to feed in the other colour. They were then tested to see whether their subsequent choices were influenced by this observation. For this, demonstrators in the previous task were separated in two groups: one group remained the “demonstrators” while the other became the new “observers”. Two individuals which had learnt to choose different colours were housed together in a tank and left to acclimate for two days. Then, as before, the new observers could first observe a demonstration during four sessions, at the end of which they were fed with the neutral stage 3 white shell to avoid reinforcement of the previously learnt colour. Second, during four sessions, they could observe the demonstrator and then were presented with the two-coloured shell apparatus five times (see above). If fishes show social learning, then the observers should reverse their preference for one colour and go feed more frequently in the other colour which has been demonstrated.

### Statistical analyses

All statistical analyses were conducted using R (version 4.2.2). For every associative learning and social learning experiment, two-tailed exact binomial tests were conducted to investigate whether a fish preferentially chose one coloured shell or one location of the shell (to the left or right of the fish home shell) (null hypothesis set at 50 %, “stats” package in R).

In the social learning experiment with naïve observers, to explore behavioural differences between demonstrators and observers when they were first presented to the experimental apparatus, we compared decision times in the first session of presentations. Decision times were defined as the time between the apparatus touching the ground and the fish’s head passing the threshold of one of the coloured shells. We ran a generalised linear mixed effect model with a negative binomial family with “role” (demonstrator or observer) as explanatory variable and “individuals” as random effect (package “glmmTMB” in R; Brooks et al., 2017). We analysed the model residuals thanks to the DHARMa package (Hartig, 2017). We then ran a type II ANOVA to analyse the contribution of our factor of interest in the variance (package “car” in R; Fox & Weisberg, 2019). Note that results should be taken with care due to the paucity of the data.

## 3. Results

### Training

The engagement and success in the training steps were low. Only one *N. multifasciatus* succeeded stage 1, learning to eat brine shrimp from a tube connected to a syringe. Other *N. multifasciatus* individuals that failed to engage with any apparatus never left their home shells and fed from drifting shrimp. In the other species, three *L. ornatipinnis* (Charlie, Bowey-Dee, and Frank) and two *L. ocellatus* (Mac and Dennis) reached stage 3 and learned to eat from a complete neutral white shell (**Table I**). Charlie and Frank (*L. Ornatipinnis*), as well as Mac and Dennis (*L. ocellatus*), were housed as pairs and frequently fed together from the stage 3 white shell during the training process. Other unsuccessful *L. ornatipinnis* and *L. ocellatus* individuals stayed at their shell and did not explore the apparatus or did not eat from it would they approach it.

**Table I.**
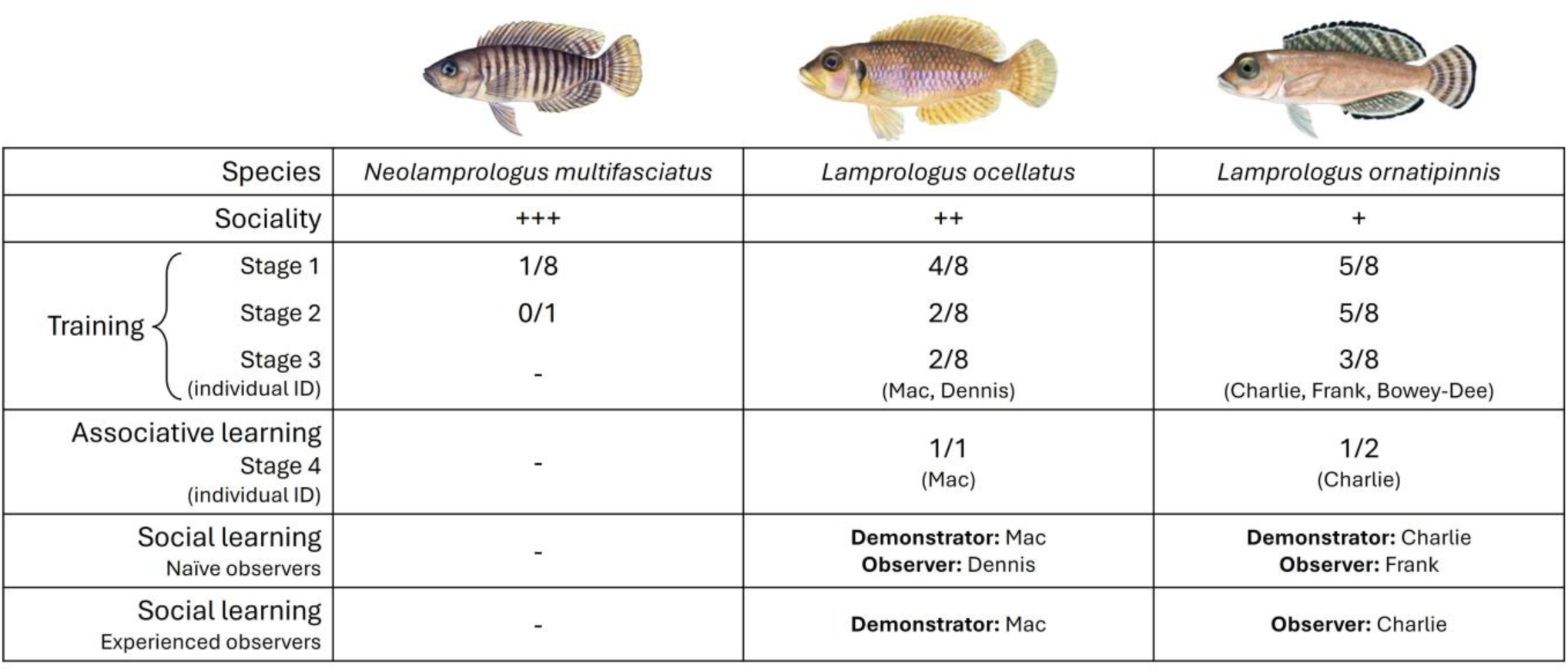
Description and results of training and experiments in the three species. Sociality signs indicate levels of sociality for each species. Numbers indicate the number of successful individuals compared to the total number of individuals for each stage of the training or associative learning experiment. Demonstrator and observers’ identities for the social learning experiments are indicated. Illustrations by Alexandra Viertler, courtesy of Jordan Lab.

For *L. ornatipinnis*, Charlie and Bowey-Dee were further tested in the associative learning task as potential future demonstrators and Frank was chosen as an observer. For *L. ocellatus*, Mac was tested in the associative learning task as potential future demonstrator and Dennis was chosen as an observer.

### Associative learning for potential demonstrators

For *L. ornatipinnis*, Bowey-Dee never engaged with the two-coloured shell apparatus and therefore did not complete the associative learning task. Only Charlie participated in this task and learned to eat from the orange shell. He chose the orange shell significantly more than the blue shell throughout all 17 sessions (ten sessions of the task alone and seven sessions while being observed by Frank) (two-tailed exact binomial test, number of successes: 76, number of trials: 96, p < 0.01) and showed no preference for a specific location of the shell (two-tailed exact binomial test, number of times choosing the shell on the right of his home shell: 52, number of trials: 96, p = 0.48; **Figure 2**).

**Figure 2:**
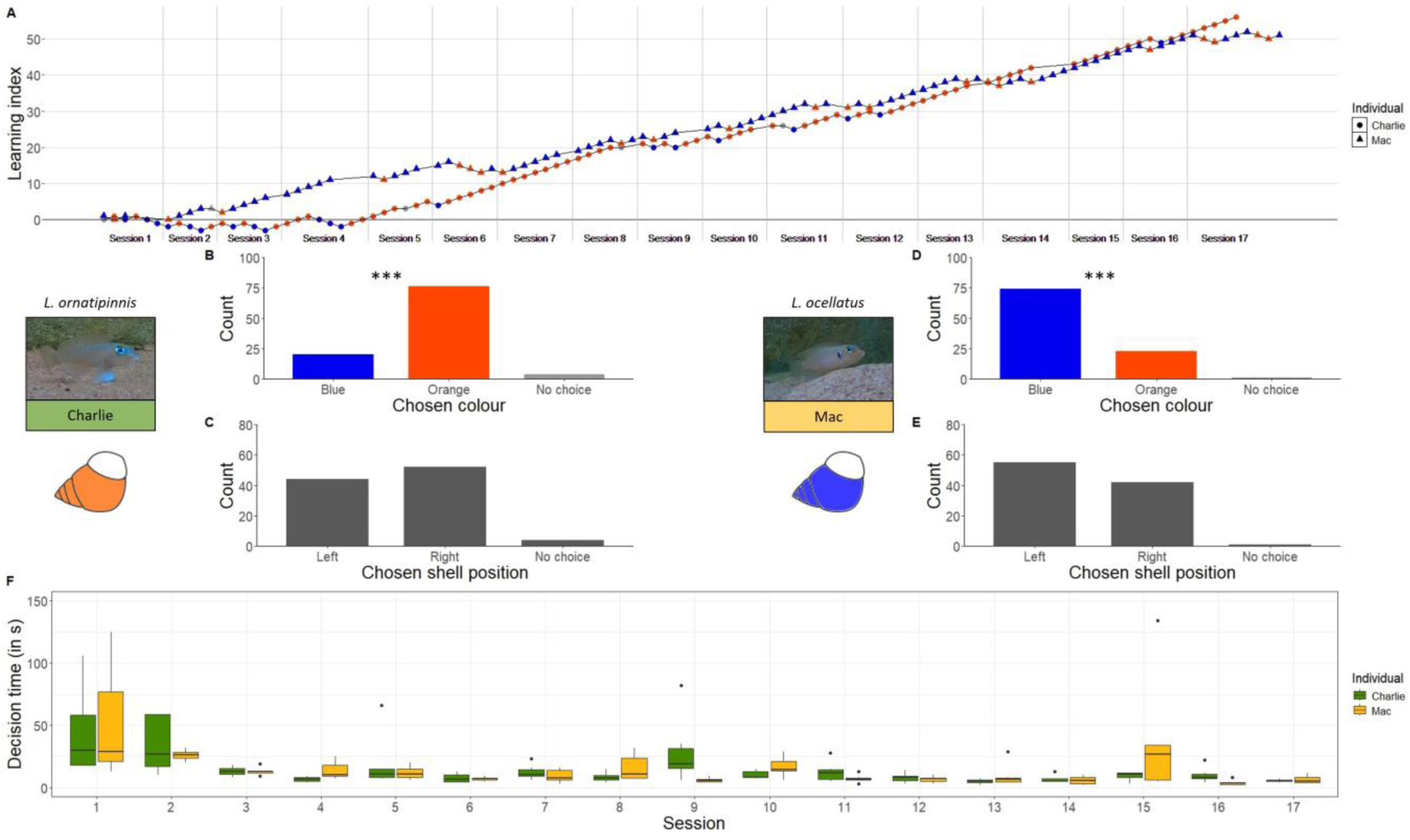
Performance of *L. ornatipinnis* (Charlie) and *L. ocellatus* (Mac) during the associative learning task. A. Learning curves of the two individuals throughout the sessions. The learning index was +1 if the fish went to the correct shell, −1 if he went to the incorrect shell, and 0 if the fish made no choice at all. From session 11 on, individuals were observed by the observers. B. Choices of the shell colour by *L. ornatipinnis* throughout all sessions. C. Choices of the shell location (right or left of home shell) by *L. ornatipinnis* throughout all sessions. D. Choices of the shell colour by *L. ocellatus* throughout all sessions. E. Choices of the shell location (right or left of home shell) by *L. ocellatus* throughout all sessions. (***: p-value < 0.001). F. Progression of decision times across sessions.

For *L. ocellatus*, Mac learned to eat from the blue shell. He chose the blue shell significantly more than the orange shell throughout all 17 sessions (two-tailed exact binomial test, number of successes: 74, number of trials: 97, p < 0.01) and showed no preference for a specific location of the shell (two-tailed exact binomial test, number of times choosing the shell on the right of his home shell: 42, number of trials: 97, p = 0.22; **Figure 2**).

Both fishes showed a decrease in their decision times across sessions and already took quick decisions from the third session on (often less than 20s) (**Figure 2F**).

### Social learning in naïve observers

For *L. ornatipinnis*, after observing Charlie eating from the orange shell, Frank did not show any preference for one coloured shell or the other (two-tailed exact binomial test, number of choices of the orange shell: 12, number of trials: 23, p = 1). However, he showed a strong location bias and chose significantly more to feed from the shell on the right of his home shell than on the left (two-tailed exact binomial test, number of times choosing the shell on the right of his home shell: 19, number of trials: 23, p < 0.01; **Figure 3**).

**Figure 3:**
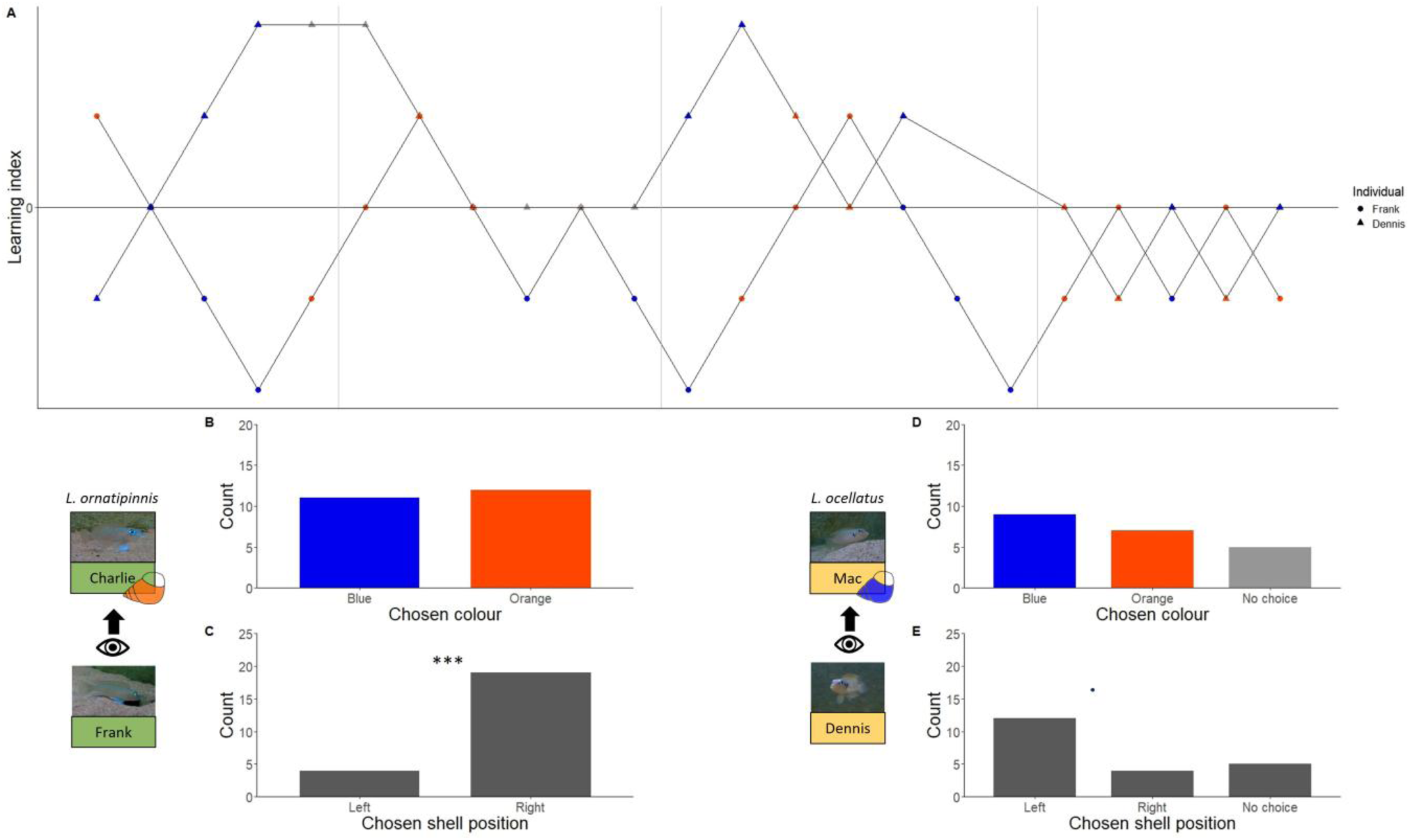
Performance of *L. ornatipinnis* (Frank) and *L. ocellatus* (Dennis) in the social learning task after having observed demonstrators. A. Learning curves of the two individuals throughout the sessions. The learning index is +1 if the fish went to the demonstrated shell, −1 if the fish went to the other shell, and 0 if the fish made no choice at all. B. Choices of the shell colour by *L. ornatipinnis* throughout all sessions. C. Choices of the shell location (right or left of home shell) by *L. ornatipinnis* throughout all sessions. D. Choices of the shell colour by *L. ocellatus* throughout all sessions. E. Choices of the shell location (right or left of home shell) by *L. ocellatus* throughout all sessions. (*: p-value < 0.5; ***: p-value < 0.001)

For *L. ocellatus*, after observing Mac eating from the blue shell, Dennis did not show any preference for one coloured shell or the other (two-tailed exact binomial test, number of choices of the blue shell: 9, number of trials: 17, p = 0.56). He demonstrated no clear preference for a specific shell location, even though he might show a tendency to choose the left shell (two-tailed exact binomial test, number of times choosing the shell on the right of his home shell when he chose: 4, number of trials: 16, p = 0.08; **Figure 3**).

When presented with the coloured shell apparatus for the first time (first sessions), both observers took significantly less time to engage with the apparatus than the demonstrators (ANOVA χ^2^ = 32.3, p < 0.001; mean decision time for demonstrators = 110.3s; mean decision time for observers = 8.4s; **Figure 4**). A simple lack of motivation from the demonstrators during all or certain trials of their first session cannot explain this difference as the minimum decision time throughout the session was 18s for *L. ornatipinnis* and 13s for *L. ocellatus*. Moreover, demonstrators showed behaviours consistent with an interest towards the apparatus (approach, going back and forth between the apparatus and their home shell…).

**Figure 4:**
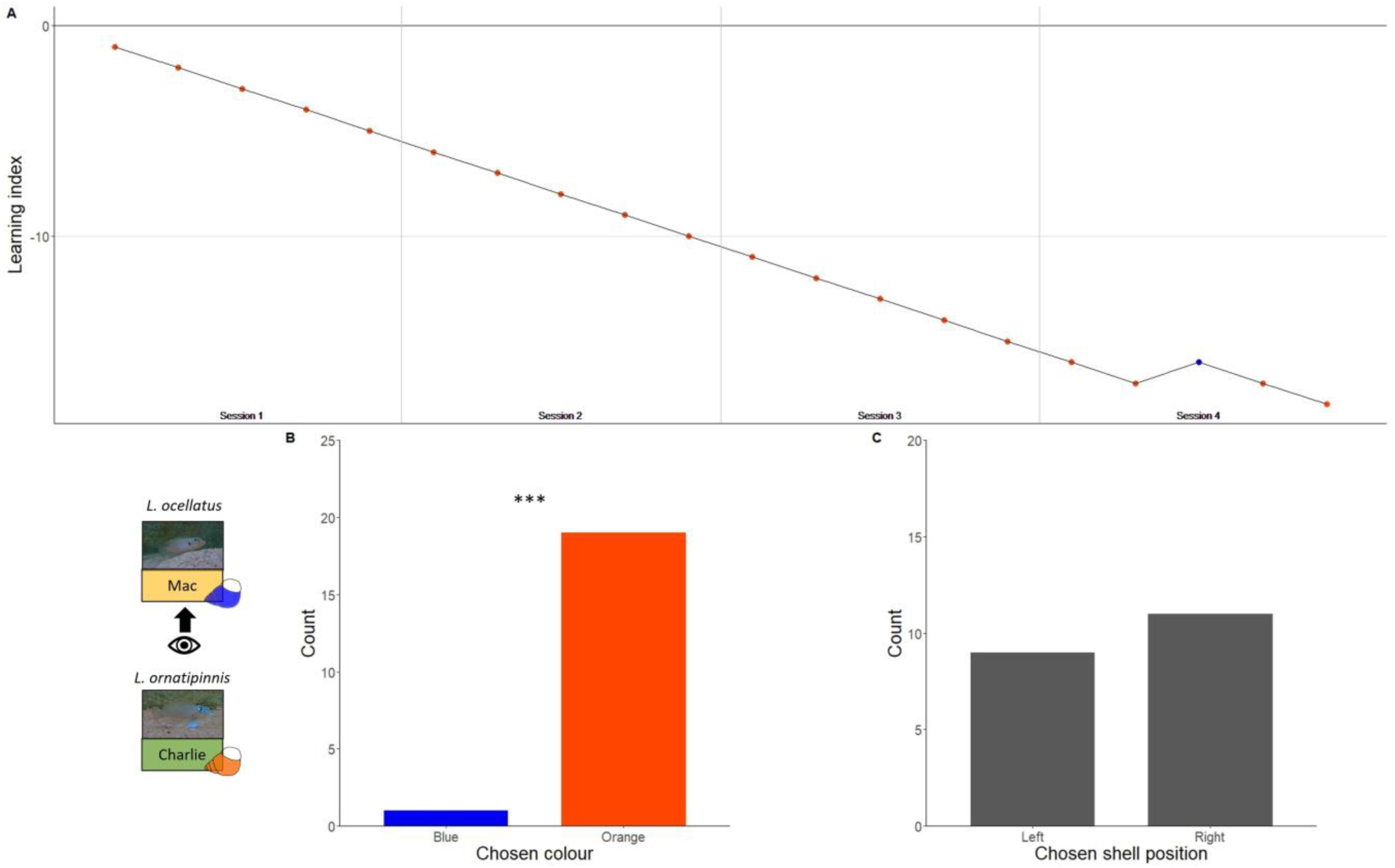
Performance of *L. ornatipinnis* (Charlie) in the social learning task after having observed *L. ocellatus* (Charlie). A. Learning curves of *L. ornatipinnis* throughout the sessions. The learning index was +1 if he went to the blue shell (demonstrated shell), −1 if he went to the orange shell, and 0 if the fish made no choice at all. The position of the chosen shell compared to the home shell is indicated below the points (L: Left, R: Right). B. Choices of the shell colour by *L. ornatipinnis* throughout all sessions. C. Choices of the shell location (right or left of home shell) by Charlie throughout all sessions. (***: p-value < 0.001)

**Figure 5:**
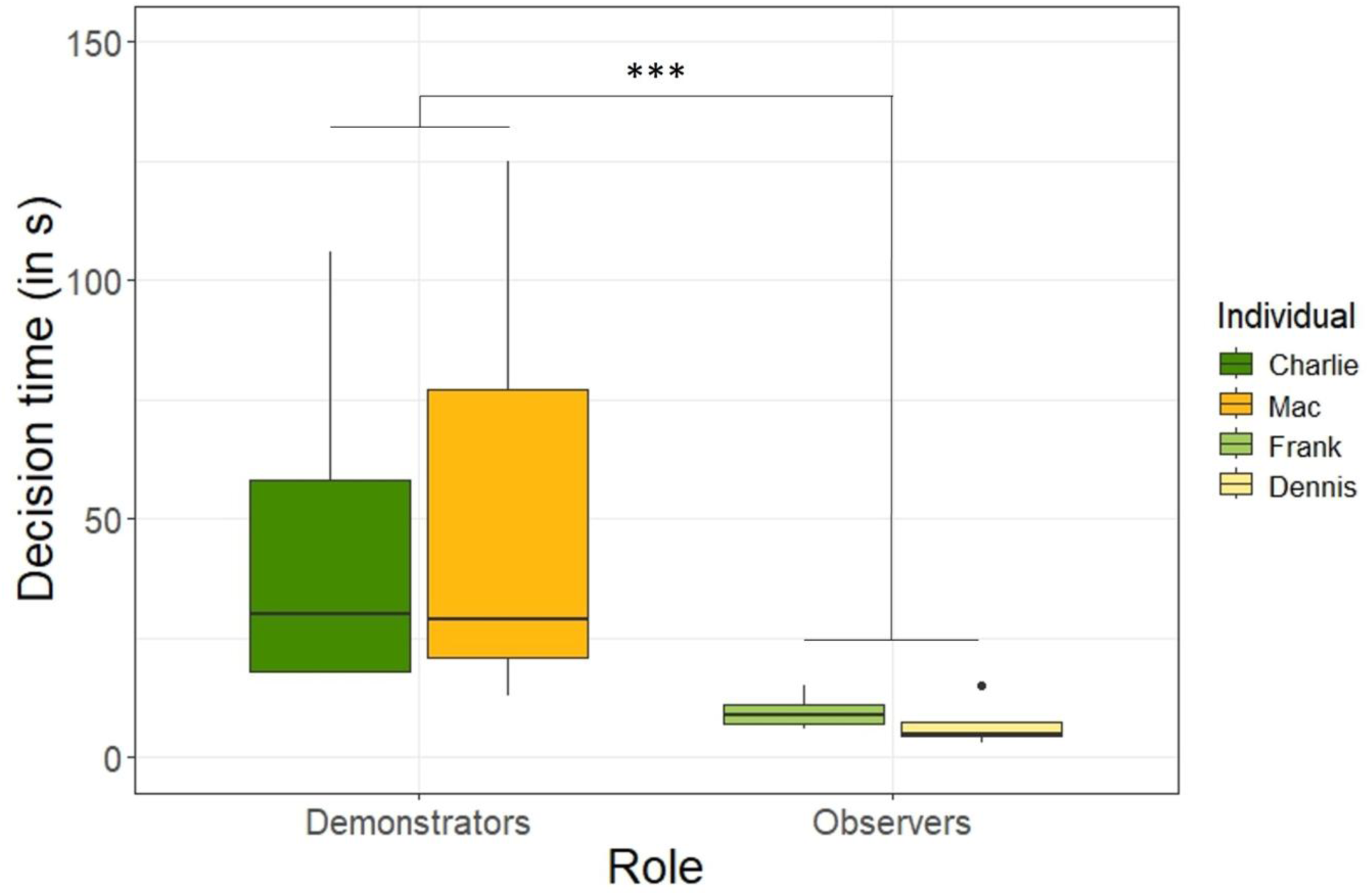
Comparison of the decision time to first feed from a shell during the first presentation of the two-coloured shell apparatus to the naïve fish (demonstrators and observers). A datapoint at 531s for *L. ornatipinnis* Charlie was removed for clarity. (***: p-value < 0.01)

### Social learning in experienced observers

Due to the low engagement in the tasks, we had only one demonstrator of both species at this stage. We thus conducted this experiment in a heterospecific context. Charlie (*L. ornatipinnis*), who had learnt to eat from the orange shell, observed Mac (*L. ocellatus*) eating from the blue shell. After the observations, in the four test sessions, Charlie did not change his training preference and almost always chose the orange shell (two-tailed exact binomial test, number of times choosing the orange shell: 19, number of trials: 20, p = 4.00.10^-5^). He did not show any preference for a specific shell location (two-tailed exact binomial test, number of times choosing the shell when on the right of his home shell: 11, number of trials: 20, p = 0.82) (**Figure 4**).

## 4. Discussion

In this study we used an associative learning paradigm in which wild Lamprologine shell-dwelling cichlids had to learn to go to a specific coloured shell to obtain a food reward. Of the three species tested, one species did not engage at all with any training step (*Neolamprologus multifasciatus*). One individual each of *Lamprologus ocellatus* and *Lamprologus ornatipinnis* successfully learnt the colour association. In the subsequent social learning tasks, none of the three individuals that learnt to feed from neutral shells (one *L. ocellatus*, two *L. ornatipinnis*), nor those that had learnt to feed from a specific colour shell (one *L. ocellatus*, one *L. ornatipinnis*) showed evidence for social learning abilities regarding colour choice, as observers did not take into account the colour of the shell in which the demonstrators were feeding, whether the observers were naïve or had previously learnt to feed from a specific alternative colour.

Our experiment proves to be another illustration of a fundamental difficulty in comparative psychology: researchers need to constantly juggle between standardizing their paradigms and adapting them to each species to obtain comparable results (Bastos & Taylor, 2020; MacLean et al., 2012). In the hope of developing a standardized paradigm for a wide array of wild shell-dwelling cichlids, we used the same experimental design for our three species of interest that were phylogenetically close. However, *Neolamprologus multifasciatus*, the most social species did not engage with even the simplest steps of training. In a similar learning task in which fishes had to eat from a well-plate, six Lamprologine shell-dwelling species showed poor engagement, even though they were all captive-bred, including the highly social species *N. multifasciatus* of which only 5 out of 30 participated in the task (Stanbrook et al., 2020b). It may be difficult for shell-dwelling species to engage in cognitive tasks because they tend to take shelter at any potential threat. Non-cognitive processes, such as neophobia can affect cognitive performances, and can also be linked to social organisations. For example, in corvids, more social species are less neophobic than species living in pairs (Miller et al., 2022). This correlation may not be true in Lamprologine cichlids where more social species seemed more wary of the apparatuses, potentially due to increased neophobia or the more extreme change in social conditions between wild and experimental contexts. Nevertheless, some individuals showed consistent engagement with our paradigm and valuable lessons and future directions can still be drawn from their responses.

Successful individuals of *L. ocellatus* and *L. ornatipinnis* showed good performance in the associative learning task, and quickly learnt to approach the correct shell. During social learning trials, we did not find evidence for colour stimulus enhancement in naïve observers. If the focal animals had shown stimulus enhancement based on the colours, they should have gone more to the colour from which they had observed the demonstrator feed. Only one *L. ocellatus* and one *L. ornatipinnis* could be tested, and neither did so. Instead, they showed strong location bias, especially in *L. ornatipinnis*. We did not observe behaviours that suggested observers simply ignored the demonstrators during observation sessions. Rather, all observers appeared very focused on the demonstration: they were located out of their home shell, bodies oriented toward the demonstrator, and they showed motivation to exit the transparent container to approach the apparatus and the demonstrator. They therefore had every opportunity to learn from the demonstrators. Nevertheless, even if the choice of colour was not socially influenced, both fishes showed some stimulus enhancement toward the new situation or new apparatus. Indeed, when they engaged with the coloured shell apparatus for the first time, naïve observers were quicker to approach the apparatus than were demonstrators when first exposed to this apparatus. They therefore appeared to have gained social information when they observed the demonstrators feeding from this novel apparatus. An alternative explanation could be a simple visual habituation to this apparatus (they could see it for three sessions before their first presentation). However, from the first session of observation already, observers showed motivation to approach the apparatus and therefore the faster approach to the apparatus when presented for the first time seems to be socially driven.

We did not find evidence for colour stimulus enhancement in experienced individuals either. One *L. ornatipinnis* that had previously learnt to go feed in one colour was not influenced by the observation of one *L. ocellatus* feeding from the other colour, although it showed all signs of paying attention during the demonstrations. Due to the small number of demonstrators available, we had to conduct this experiment in a heterospecific context, and it is possible that heterospecific information was less relevant to the animals. There is however no hint that it is the case and that heterospecific social learning is rarer than conspecific social learning, even more so in relatively solitary species (Avarguès-Weber et al., 2013; Seppänen et al., 2007; Webster & Laland, 2017). Cichlids living in syntopy, there are great chances that they are as fit to learn from heterospecifics than conspecifics (Lein & Jordan, 2021).

Nevertheless, it seems that the observers prioritised private information against public information in their choices of colours. Multiple species show situations where they prefer private over public information (*e.g.*, honeybees (Grüter et al., 2013), minnows (Webster & Laland, 2008) or ants (Stroeymeyt et al., 2017)). Several explanations have been proposed for the prioritization of private information, such as the lack of crucial dimensions of public information (Czaczkes et al., 2019). In our paradigm, like in artificial fruits paradigms (*e.g.*, Dindo et al., 2011), we rewarded any colour chosen by the observer. Using private over public information was therefore not costly and may have encouraged individuals to rely on private information regarding the colours. However, when first presented with a new apparatus, naïve observers appeared to use public information and approached the apparatus quicker than the demonstrators. Moreover, for both species, the individuals that succeeded in all the training steps were housed in the same tank, suggesting social facilitation effects. Both species therefore seemed to gain information over novel objects socially. Future studies could further explore the impact of social information on neophobia in shell-dwelling cichlids, using observational avoidance conditioning tasks like Barks & Godin, 2013.

Based on the social intelligence hypothesis, we hypothesized that more social species would show better associative learning and social learning abilities. Due to the lack of comparison with *N. multifasciatus*, we cannot draw any firm conclusion on species differences that could help shed light on the merits of this hypothesis in our studied species but both *L. ornatipinnis* and *L. ocellatus* showed quick associative learning abilities. Being relatively non-social species, it is not surprising that they would tend to base their decisions on private rather than public information in the social learning trials. Neither species showed major differences in their learning processes, but our sample size impedes us from drawing firm conclusions. Given their incredible diversity in socio-ecological factors, cichlids are a model of choice to test cognitive and non-cognitive processes and our findings, despite low engagement from wild fishes, will hopefully invigorate further investigations of social learning abilities in this group.

## 5. Acknowledgements

The authors gratefully acknowledge their collaborators at the Department of Fisheries in Mpulungu, Zambia, and at the University of Zambia.

## 6. Declarations

### Ethical approval

We declare that our study is in full accordance with the ethical guidelines of our institutions and comply with the Zambian and European legislation on animal welfare. The experiments were conducted according to the Guidelines for the treatment of animals in behavioural research and teaching (https://doi.org/10.1016/S0003-3472(21)00389-4) and were conducted under Zambian Study Permit numbers SP268511/9-21 and SP297185/4-22 and in conjunction with a memorandum of understanding with the University of Zambia (MOU 101/14/11). The study species are listed as ‘Least Concern’ on the IUCN Red List of Threatened Species.

### Author’s contributions

M.T., V.D., and A.J. conceptualised the experiments. M.T. and M.S. performed the experiments. M.T. analysed the data, performed the analyses, prepared the figures, and wrote the main manuscript text. V.D. and A.J. reviewed the manuscript. V.D. and A.J. contributed equally in supervising this work.

### Availability of data and materials

Data and R script are available on request to the corresponding author.

